# Capturing scientific knowledge in computable form

**DOI:** 10.1101/2021.03.10.382333

**Authors:** Jeffrey V. Wong, Max Franz, Metin Can Siper, Dylan Fong, Funda Durupinar, Christian Dallago, Augustin Luna, John Giorgi, Igor Rodchenkov, Özgün Babur, John A. Bachman, Benjamin M. Gyori, Emek Demir, Gary D. Bader, Chris Sander

## Abstract

Technological advances in computing provide major opportunities to accelerate scientific discovery. The wide availability of structured knowledge would allow us to take full advantage of these by enabling efficient human-computer interaction. Traditionally, biological knowledge is captured in publications and knowledge bases, however, the information in articles is not directly accessible to computers, and knowledge bases are constrained by finite resources available for manual curation. To accelerate knowledge capture and communication and to keep pace with the rapid growth of scientific reports, we developed the Biofactoid (biofactoid.org) software suite, which crowdsources structured knowledge in articles from authors. Biofactoid is a web-based system that lets scientists draw a network of interactions between genes, their products, and chemical compounds and employs smart-automation to translate user input into a structured language using the expressive power of a formal ontology. The resulting data is shared via public information resources, enabling author-curated knowledge to be appreciated in the context of all existing computable knowledge. Authors of recently published papers across a range of journals have already contributed their pathway information, much of which is novel and extends existing pathway databases into new biological areas. We envision the adoption of Biofactoid for crowdsourced curation by scientists and publishers as part of an ecosystem of tools that accelerate scientific communication and discovery.

**Availability:** Biofactoid server at https://biofactoid.org

## INTRODUCTION

Biological pathways organize sets of molecular interactions and reactions that underlie cellular processes and are used for experimental design, interpretation of genomics data (Khatri et al., 2012), understanding of disease mechanisms (Chinen et al., 2016; Santos et al., 2014), and identification of therapeutic targets (Mack et al., 2014). To manage, visualize and interpret the large amount of available pathway information, researchers require computational tools such as pathway information resources, structured data representation standards, and analysis software (Demir et al., 2010; Franz et al., 2016; Jassal et al., 2020; Pratt et al., 2015; Rodchenkov et al., 2020; Shannon et al., 2003). These computational tools are capable of accessing increasing amounts of pathway and interaction data (Bader et al., 2006) collected through centralized (Gene Ontology Consortium, 2015; Jassal et al., 2020) or crowdsourced curation efforts (Slenter et al., 2018). Nevertheless, these existing efforts are not able to achieve wide coverage of the rapidly growing scientific corpus (over 1.3 million new PubMed articles/year) (Bornmann and Mutz, 2015; Cordero et al., 2016), arguing for the development of more scalable and sustainable approaches (Attwood et al., 2015; Imker, 2018).

For reasons of efficiency and accuracy, structured knowledge of biomedical discoveries could be provided directly by the original authors of research reports, rather than *post-hoc*, through the curation efforts of knowledge base teams. By analogy, molecular 3D structures submitted directly by authors to the Protein DataBank (PDB) (Berman et al., 2000) have become a key resource among structural biologists, and similar community practice led to direct submission of DNA sequence and transcriptomics information to public databases. Indeed, the importance and value of having such research outputs available in a community resource are underscored by the fact that data deposition is often a requirement of publication and funding. In contrast, there are few efforts and little technology to support direct submission by authors of biological pathway information and related knowledge in computable form.

Here we introduce Biofactoid (biofactoid.org), a web-based software system that empowers authors to capture and share structured human- and machine-readable summaries of molecular-level interactions described in their publications. Without such a curation support tool, the onus would be on authors to handle a series of complex tasks involved in converting their knowledge to a computational form and depositing it in a suitable knowledge base. To overcome this and other significant barriers to computable knowledge acquisition and sharing, we developed Biofactoid to ease pathway curation and to rapidly generate expressive, structured representations with minimal user training. Structured knowledge newly acquired in this way becomes part of the global pool of pathway knowledge and can be shared in resources such as Pathway Commons (Rodchenkov et al., 2020), Network Data Exchange (NDEx) (Pratt et al., 2015), and STRING (Szklarczyk et al., 2017), to enhance information discovery and analysis. Authors can use Biofactoid to share structured information from research articles both as part of the publication process and outside of it. The development of Biofactoid and related computational tools helps support human-computer communication and inference algorithms in an ecosystem in which scientific reasoning is increasingly assisted by broad and deep knowledge computation.

## RESULTS

### Data sharing workflow

Biofactoid enables molecular-level detail of biological processes reported in articles to be shared in a structured format accessible to humans and computers. Interactions (including binding, post-translational modification, and transcription/translation) involving molecules of various types (proteins, nucleic acids, genes, or chemicals, e.g., metabolites and drug compounds) can be represented. Users begin by entering the article title, or identifier (e.g. PubMed identifier or Digital Object Identifier). Article metadata including authors, abstract and journal issue is automatically retrieved. Next, users draw a network of biological entities and interactions using the Biofactoid curation tool, which has an easy-to-use interface similar to graphical illustration software (e.g. Microsoft Powerpoint, Adobe Illustrator) which will be familiar to users, but with the added ability to generate structured data from the author-drawn biological pathway. The major features of the Biofactoid curation tool are as follows:

1. **Molecular entities**. Genes, gene products and chemicals are created in the network using an “Add a gene or chemical” tool, which creates a node (circle) that users label (Figure 1A). Biofactoid automatically matches the label with a record in an external database: NCBI Gene for genes and their products and ChEBI for small molecules (Brown et al., 2015; Hastings et al., 2016). This match represents the top hit of a search based on the similarity between a user’s label and a database record’s list of common names and synonyms; for genes, organisms are given priority based upon the organism of genes previously added to the network. Users can update the match by selecting another organism (e.g. human or mouse p53) or update the automatically inferred gene product type (i.e. RNA or protein). Alternatively, users may assign an alternate database entry from a list of search results. The system supports human and select model organisms (M. musculus, R. norvegicus, S. cerevisiae, D. melanogaster, E. coli, C. elegans, D. rerio, and A. thaliana). The system is fast (response times of 100 milliseconds or less) and can be easily extended, for instance, to support SARS-CoV and SARS-CoV-2, which we added in response to the burst of publications related to the COVID-19 global pandemic (Ostaszewski et al., 2020). The matching process is also accurate, as measured using tests that use entity names from research articles as queries and assessing the quality of the search result. In over 90% of the cases, the correct result was first among search hits and was among the top 10 in over 97% of cases. Thus, Biofactoid can offer an accurate database identifier match using only the author-provided entity name. This represents a major advance in usability compared to traditional gene and chemical name querying systems that only work with exact name string matching and are often slow or unreliable. In fact, we were forced to build our own system after trying to use major existing systems that turned out to have high failure rates for our use case.

**Figure 1.**
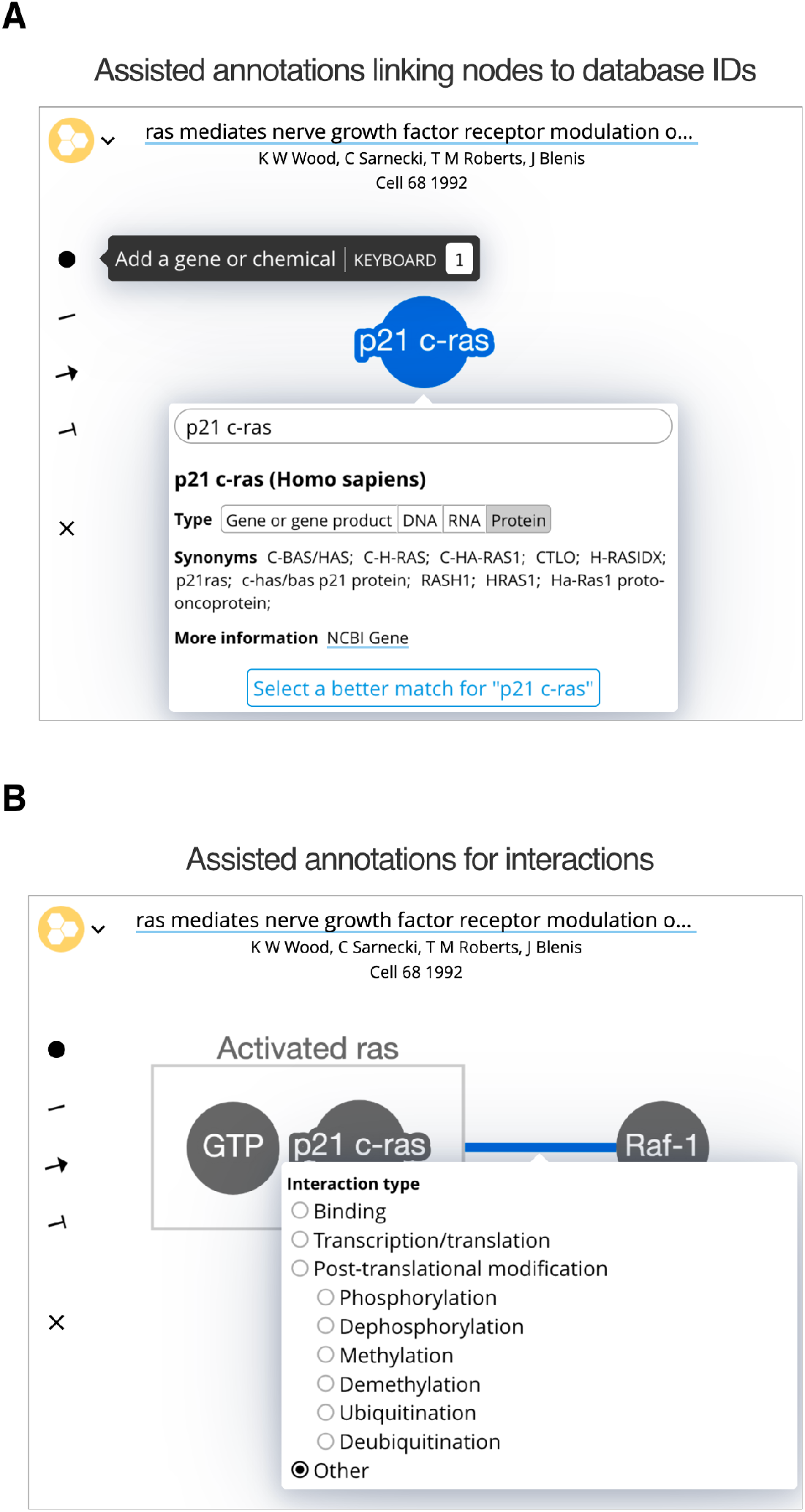
The Biofactoid curation tool. Curation in Biofactoid involves drawing a network of relationships between genes or molecules. **(A)** Genes and chemicals are represented by circles (nodes) where users provide a label, the type of gene product, and the organism. A custom search engine matches the label to a corresponding record from a database of genes or chemicals. **(B)** Relationships are represented by connecting participants with lines, arrows (to indicate activation), or ‘T-bars’ (to represent repression). Users select the mechanism that best describes the interaction. Complexes are represented as participants enclosed by a box.
2. **Interactions**. A “draw interaction” tool lets users link two nodes by clicking one and dragging to the other (Figure 1B). Edges can have type and direction: a pointed arrow indicates stimulation or activation; a ‘T-bar’ arrowhead indicates inhibition or repression; and an undirected line indicates any other interaction. The mechanism can be refined by selecting from a list that includes binding, transcription/translation, or common forms of post-translational modifications, with additional interaction types to be added in future releases.
3. **Complexes**. Molecular complexes can be created by dragging genes or chemicals close to each other, resulting in a box that encloses them, which can also be labeled (Figure 1B).
4. **Automatic saving and co-editing**. Pathway representations in Biofactoid are automatically saved as changes are made and a live-sync capability enables multiple authors to collaboratively edit the same pathway, analogous to Google Docs.

Once complete, the pathway data is validated by pressing a ‘Submit’ button. At this point, the user may address any potential quality issues flagged by the system (e.g. unlabelled nodes, empty document) before confirming their submission.

### Data sharing and exploration

When research findings are shared through Biofactoid in a structured and computable manner, curated knowledge is automatically connected to, and becomes part of, a collective pool of computable knowledge that the community can access in different ways. Visitors to the Biofactoid website (biofactoid.org) may browse recently added articles, and a graphical abstract of each new submission is posted to Twitter (twitter.com/biofactoid). Each entry is automatically linked to its associated authors, article information and structured data and presented in an interactive Biofactoid Explorer web app (Figure 2A). Users can select any entity in the pathway diagram to see more information about genes (via NCBI Gene database links), proteins (via UniProt database links (UniProt Consortium, 2019)) and chemicals (via ChEBI database links). To support attribution, each author of an article is automatically linked to their profile in the Open Researcher and Contributor ID database (ORCID; orcid.org). Finally, a set of ‘Recommended articles’ are generated to help the user explore similar knowledge in the literature. To generate these recommendations, we first retrieve an article’s references, articles it is cited by and “Similar articles’’ from PubMed using the NCBI Entrez Programming Utilities service (Wheeler et al., 2006). These are combined with articles that mention or provide evidence for interactions involving any of the molecular entities in the Biofactoid entry, which are accessed from curated pathway and interaction databases as well as resources that aggregate interaction information directly from the biomedical literature using natural language processing (NLP) tools (e.g. REACH, INDRA) (Gyori et al., 2017; Valenzuela-Escárcega et al., 2018). Recommended articles are ranked by determining how similar their titles and abstracts are to that of the Biofactoid article, using a deep-learning-based method we developed (Giorgi et al., 2020). The list of recommendations is context-sensitive in that articles are shown relevant to the part of the pathway selected by a user.

**Figure 2.**
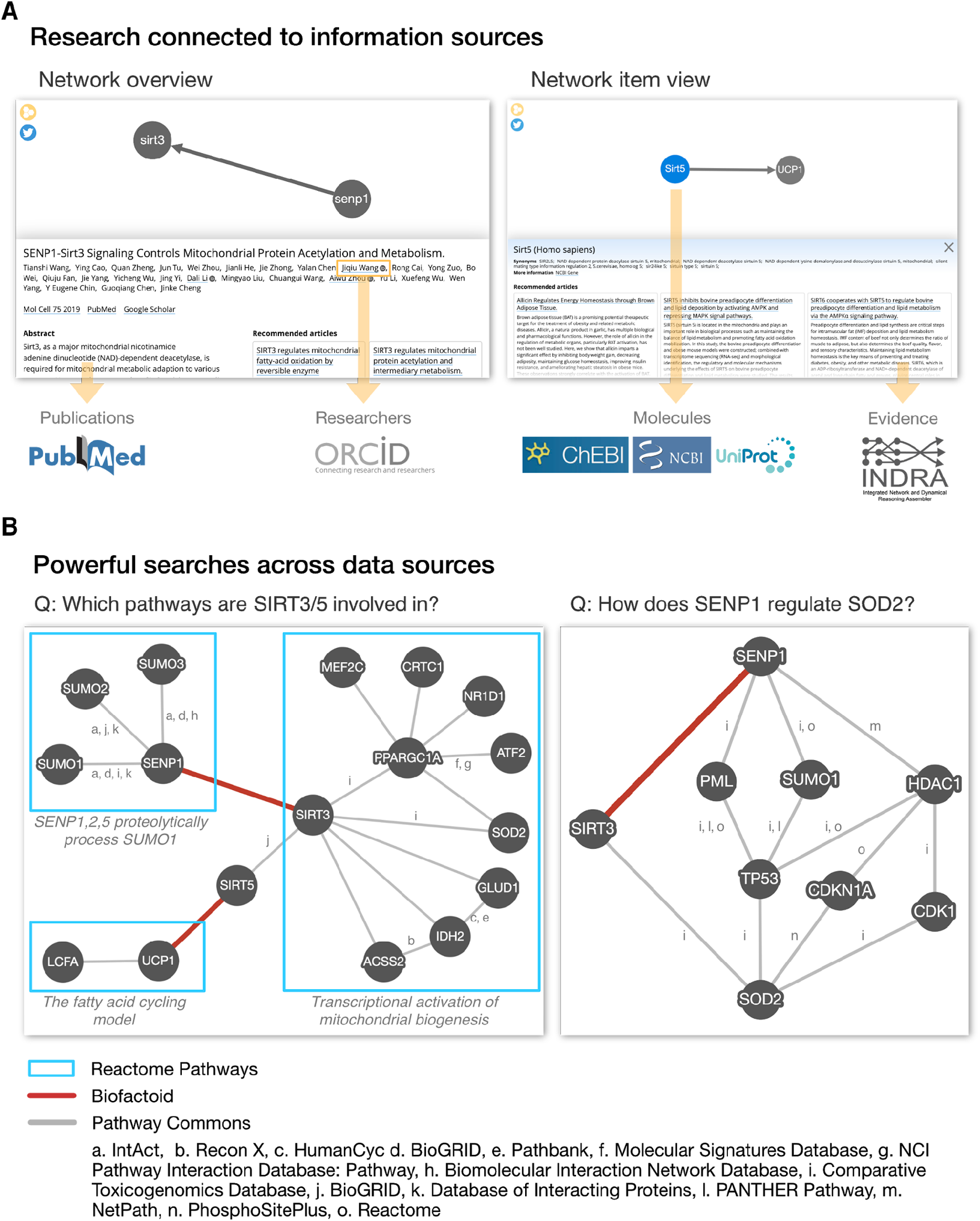
Biofactoid data is connected to existing knowledge and enables more powerful search. **(A)** The Biofactoid Explorer is an interactive web app that publicly presents each author-curated entry alongside their article. Arrows indicate how curated information is connected to outside knowledge bases. A “Network overview” (left) displays information about the article and pathway as a whole; a “Network item view” (right) displays information for a selected item (e.g. interaction, protein). **(B)** Biofactoid data integrated with existing structured biological knowledge enables powerful searches across data sources. Two author-curated interactions submitted to Biofactoid (red edges) bridge previously distinct pathways from the Reactome Pathway Database involved in mitochondrial biogenesis (left) and provide a new, more direct regulatory route between two mitochondrial genes (right). Pathway and interaction information was provided by Pathway Commons (pathwaycommons.org), a web resource that provides a single point of access for multiple public interaction and pathway databases.

Beyond the ‘Explorer’, Biofactoid data represented in the formal BioPAX data exchange language is integrated with external pathway and interaction knowledge using technologies for BioPAX processing and analysis, including Paxtools (Demir et al., 2013) and cPath2 (Cerami et al., 2006). This enables author-contributed information to be more easily searched, visualized and analyzed across different data sources. For example, researchers can ask the question: “*Which known pathways are related to an interaction in Biofactoid?*” As an example answer to this question, a search of Pathway Commons shows how two interactions made computable by authors using Biofactoid (i.e. “*SENP1 activates SIRT3”* (T. Wang et al., 2019); “*SIRT5 activates UCP1*” (G. Wang et al., 2019)) unify previously disconnected pathways for mitochondrial biogenesis from the Reactome Pathway Database (Jassal et al., 2020) (Figure 2B). Researchers can also ask: “*How are two genes related?*” As another query example, a search of Pathway Commons shows how an interaction contributed by an author using Biofactoid (“*SENP1 activates SIRT3”* (T. Wang et al., 2019)) provides a new, more direct regulatory pathway linking the mitochondrial proteins SENP1 and SOD2. Thus, Biofactoid directly contributes to building a more complete and comprehensive collection of biological pathways.

### Pilot study involving authors and journals

We tested the Biofactoid software over many iterations of testing involving authors and journal editors, extensively updating the software each time based on user feedback. This process supported two major goals: to improve the user experience of the software for authors, who are typically unfamiliar with curation and structured data concepts, and to develop a model for integrating Biofactoid into the publication process with journals. Once Biofactoid software achieved a sufficient level of sophistication and completeness, satisfying many of these initial users, we engaged journal editors to determine the feasibility of using Biofactoid to capture information from authors. Our pilot study consisted of three phases (Figure 3). Phase I introduced Biofactoid to journal editors via an “author-simulation”, which began with a mock email invitation asking the editor to use Biofactoid to curate a selected article, and ending when a pathway from that article was input into the system by the editor using the Biofactoid web-based user interface. Phase II involved sending email invitations to a small number (N=15) of authors from a single journal, with multiple successful responses, proving that authors can use the system without any direct support from the Biofactoid team or journal. Phase III measured engagement rate by emailing 260 authors who published selected articles across 16 journals. Roughly 8.5% of these authors successfully shared their research in Biofactoid, proving that many authors can and will use Biofactoid (Figure 2A). We also found that 10% of articles across selected journals are suitable for inclusion in Biofactoid based on the current set of supported concepts (Table S1). In addition to proving that Biofactoid can be independently used by authors to capture pathway knowledge, these results indicate a strong willingness among authors in the research community to use the system. These also demonstrate that the information captured by authors may not be present in any existing pathway database and helps connect previously unconnected entities and pathways in these databases (Figure 2B).

**Figure 3.**
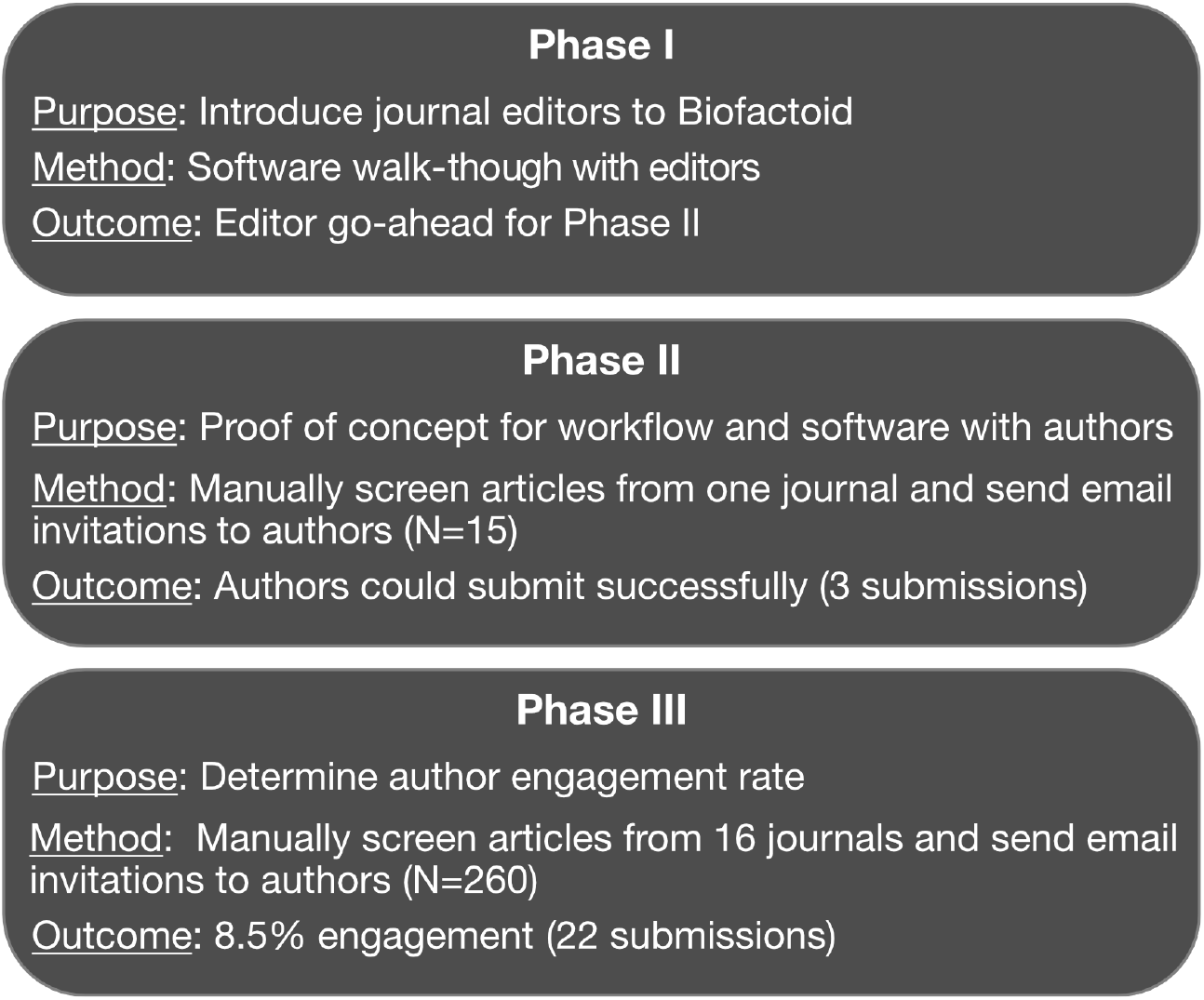
Biofactoid pilot study. A three-phase pilot tested the feasibility of Biofactoid and involved journal editors and authors of research articles. Phase I and II involved editors and authors whose articles were recently published. In Phase III, 2065 published articles were screened and authors of suitable articles were invited to Biofactoid. The articles screened were from 16 journals (Table S1).

## DISCUSSION

Biofactoid focuses on the capture of published pathway information in computational form to augment discovery, attribution, and communication of scientific knowledge. In developing our strategy, we have considered how technology can best be used to aid authors. The result is a generic approach that can be extended to capture other types of biological knowledge, at the source of knowledge creation, in computable form. To be successful, Biofactoid development has focused on providing key elements of efficiency, incentives, and better technology for human-computer interaction. The computable knowledge capture model proposed here includes formal knowledge representation using ontologies, easy to use curation support software that links molecules to corresponding database identifiers (normalization), submission to widely available knowledge resources, a connection with the publication process and author attribution. This model expands on prior work defining digital abstracts and building crowdsourcing efforts for pathway data and software (Bharadwaj et al., 2017; Ceol et al., 2008; Gerstein et al., 2007; Liechti et al., 2017; Pratt et al., 2015; Slenter et al., 2018; Todorov et al., 2019; Waagmeester et al., 2020). Future development will include expanding the Biofactoid process to capture increasing amounts of literature-described pathway information such as additional relationship types, context and direct vs. indirect interactions, prioritized by frequency of occurrence in publications. In principle, Biofactoid technology can be extended to support any network of concepts and their relationships (i.e. knowledge graph) collaboratively built by a community of knowledge generators.

A convenient time to capture knowledge is during publication when authors – the primary source of knowledge - are most aware of the details of their report and when they are typically asked to deposit other types of data (e.g. sequencing or expression data). Thus, Biofactoid can integrate with the publication process, but also can be used in crowdsourcing efforts, which have been especially useful in curating knowledge about SARS-CoV-2 (Ostaszewski et al., 2020). As a demonstration, our pilot study engaged publishers to establish requirements for using Biofactoid within their publication pipeline, ideally as a condition for acceptance, and to drive greater awareness of the system among the wider readership. The pilot study also revealed that directly inviting authors of suitable articles to contribute, even after publication, is a viable engagement strategy. However, the time-consuming and labour-intensive nature of manually screening articles argues for the development of systems (e.g. using NLP) to automate article triage. If successful, this approach could be used to regularly identify relevant articles from the pool of all new entries indexed by PubMed. Another approach to reach users is to notify those whose research articles are referenced by information curated in Biofactoid. For this purpose, we have developed a system that automatically notifies authors when their research is linked to papers curated in Biofactoid (e.g. by citations or because of related content) so that they may further explore this information and curate their own pathway knowledge. More generally, we intend on engaging the research community by cultivating partnerships with knowledge database organizations including those that curate information about articles (PubMed), biomolecules (e.g. UniProt, NCBI, ChEBI), pathways and interactions (e.g. Reactome, STRING), model organisms (e.g. Saccharomyces Genome Database (Lang et al., 2018)), and researchers (e.g. ORCID).

To improve the efficiency and utility of Biofactoid, we are developing machine-learning-based NLP technology to support authors in using Biofactoid, as well as to enable the representation of pathways in textual form (Giorgi et al., 2019; Giorgi and Bader, 2020, 2018; Valenzuela-Escárcega et al., 2018). To better accommodate the way individual users prefer to communicate, the system will accept both graphical and textual entry of pathway information as well as automated conversion between these two forms. Assistants will support users to rapidly compose their network, enabling them to add new information from a list of interactions identified in their article by NLP technology (Gyori et al., 2017). Further development of NLP methods for the reliable extraction of pathway information from the publication full-text, combined with development of new tools for curation and quality control, will help realize broad and accurate coverage of pathway knowledge in computable form. We are also improving the search and exploration functions of the system, ensuring Biofactoid information is well connected to other useful knowledge, and that related information is easily accessible starting from a Biofactoid entry. In the future, we envision that information entered by an author in Biofactoid serves as a custom query that can be used to regularly notify the author of new information (e.g. from other publications) related to their interests, such as interacting molecules and phenotypes. This work provides a basis for the development of new technologies to make scientific knowledge more computable and accessible and help researchers identify information within the rapidly growing scientific corpus.

## MATERIALS AND METHODS

### Implementation

Biofactoid is written in JavaScript. The backend server uses a microservice architecture, with Node.js, Express, and RethinkDB. Client-server data synchronization, supporting automatic saving and concurrent editing, uses websockets and a model similar to differential synchronization (Fraser, 2009). The front end uses React and Cytoscape.js (Franz et al., 2016), for network drawing and is optimized for desktop and mobile devices. An administrative dashboard, as well as user curation workflow automation features (e.g. automatic email generation) are integrated into the Biofactoid system to aid system scalability.

### Availability

Biofactoid is available to biomedical researchers for data sharing and exploration on the web at biofactoid.org. To support bioinformaticians and software developers, all user-contributed pathway data is openly accessible in multiple standard formats: JavaScript Object Notation (JSON) for raw data; Systems Biology Graphical Notation Markup Language (SBGN-ML) pathway visualization format using the Process Description notation (Le Novère et al., 2009; van Iersel et al., 2012); and BioPAX (Demir et al., 2010). All code, documentation and data are open source and freely available through GitHub (github.com/PathwayCommons/factoid); containerized components are freely available on DockerHub (hub.docker.com/r/pathwaycommons/factoid) enabling others to build on and improve the Biofactoid software.

## FUNDING

Biofactoid development was funded by the US National Institutes of Health (NIH) [U41 HG006623, U41 HG003751, R01 HG009979 and P41 GM103504] and the DARPA Big Mechanism and Communicating with Computers programs [ARO W911NF-14-C-0119, W911NF-15-1-0544].

## ACKNOWLEDGEMENTS

We thank Quincey Justman, Miao-Chih Tsai, and Anita DeWaard for feedback on making Biofactoid useful for editors and authors; the Reactome database team for support and feedback on the curation workflow; Alfonso Valencia, and Miguel Vazquez for early support; and the many community members in Toronto, Boston, Portland, and beyond for feedback on the design and concept of Biofactoid.

## COMPETING INTERESTS

None declared.

## SUPPLEMENTARY FIGURES AND TABLES

**Table S1.** Prevalence of articles with pathway knowledge suitable for Biofactoid

| ISSN | <sup>1</sup>Journal | <sup>2</sup>Coverage [Vol(Iss)] | Articles screened | Hits | % Hits |
| --- | --- | --- | --- | --- | --- |
| 2211-1247 | Cell Reports | 30(1) - 32(11) | 953 | 109 | 10.3 |
| 1097-4164 | Molecular Cell | 73(1) - 79(6) | 725 | 85 | 10.5 |
| 1549-5477 | Genes & Development | 34(1-2) - 34(17-18) | 93 | 15 | 13.9 |
| 1476-4679 | Nature Cell Biology | 22(4) - 22(9) | 84 | 10 | 10.6 |
| 1083-351X | Journal of Biological Chemistry | 295(31) - 295(37) | 210 | 21 | 9.1 |
| **Total** | - | - | 2065 | 240 | - |
| **Weighted Average** | - | - | - | - | 10.4 |

<sup>1</sup>Only journals in which at least 80 'Hits' were identified were included. A 'Hit' is an article that provides direct evidence for a molecular interaction that can be captured by Biofactoid. The ability of Biofactoid to capture an interaction depends upon the type of bioentities, the relationship types and organisms described in the article. In total, articles from 16 journals were screened: EMBO; Molecular and Cellular Biology; Cell; Cancer Cell; iScience; Cell Metabolism; Science; Nature Genetics; Science Signaling; Science Advances; Immunity; Cell Reports; Molecular Cell; Genes & Development; Nature Cell Biology. <sup>2</sup>Coverage indicates the span of journal issues that were included. Only primary research articles from each issue were screened.

## Notes

### Competing Interest Statement

The authors have declared no competing interest.

